# openPFGE: An open source and low cost pulsed-field gel electrophoresis equipment

**DOI:** 10.1101/2020.06.22.165928

**Authors:** Diego Lagos-Susaeta, Oriana Salazar, Juan A. Asenjo

## Abstract

DNA electrophoresis is a fundamental technique in molecular biology that allows the separation of DNA molecules up to ~50 Kbp. Pulsed-field gel electrophoresis [PFGE] is a variation of the conventional DNA electrophoresis technique that allows the separation of very large DNA molecules up to ~10 Mbp. PFGE equipment is very expensive and it becomes an access barrier to many laboratories. Also, just a few privative designs of the equipment are available and it becomes difficult for the community to improve or customize their functioning. Here, we provide an open source PFGE equipment capable of the separation of DNA molecules up to, at least, ~2 Mbp and at low cost: USD$850, about 3% of the price of typical commercial equipment.

**Specifications table:** 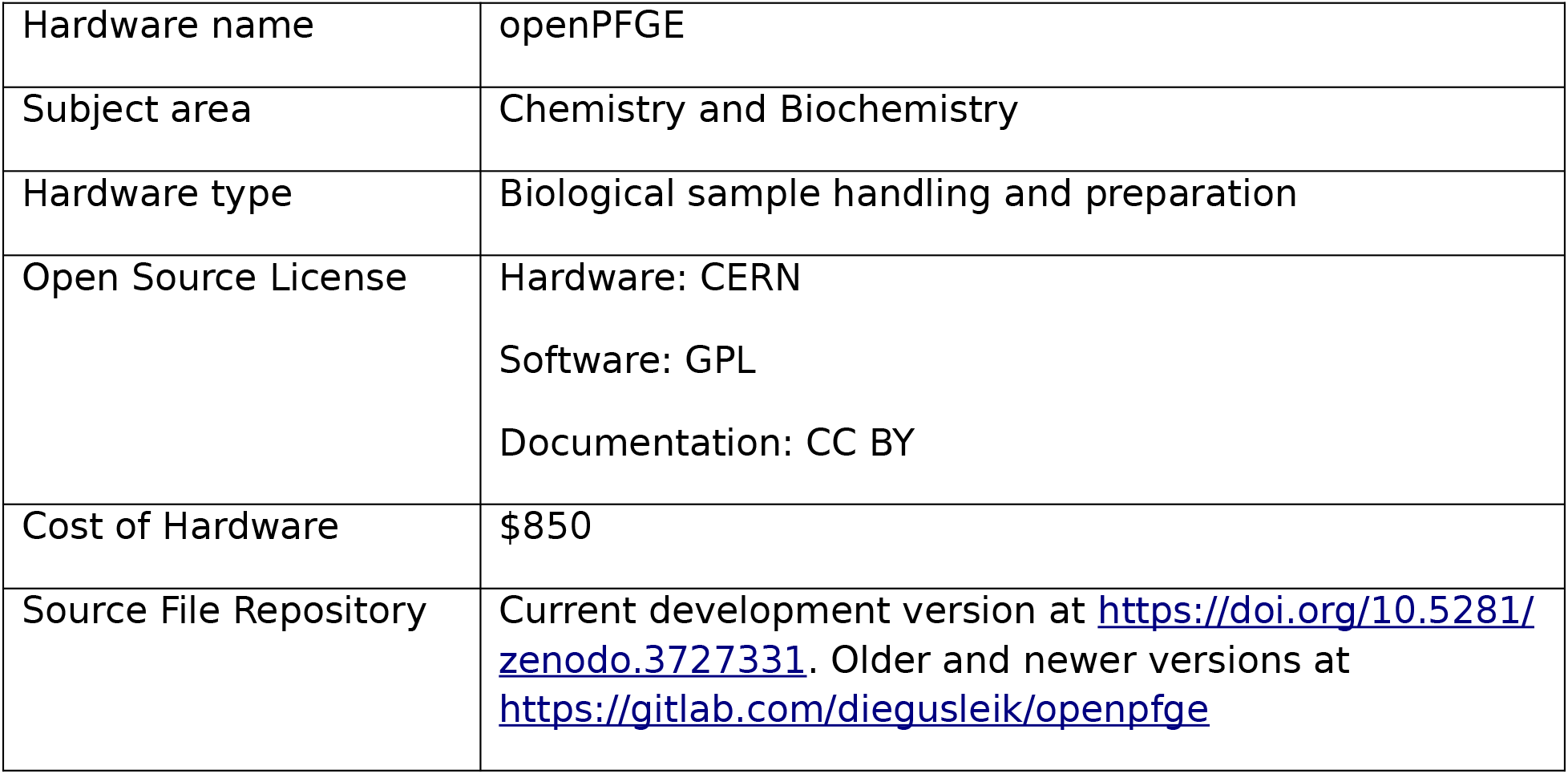

## 1. Hardware in context

DNA electrophoresis is a frequently used technique in molecular biology that allows the separation of DNA molecules up to ~50 Kbp by applying a single-direction electric field across a slab of agarose gel where small DNA molecules are sieved in a size-dependent manner. However, the sieving effect underlying such separations fails when very large DNA molecules must be resolved, as is the case of genomic analysis using intact chromosomal DNA molecules or analysis and manipulation of very large restriction fragments. Pulsed-field gel electrophoresis [PFGE], developed by Schwartz and Cantor (1), is a variation of the conventional DNA electrophoresis technique that allows the separation of very large DNA molecules up to ~10 Mbp. This is achieved by a periodic and abrupt change in the direction of the electric field that exploits the reptation phenomena of large DNA molecules to allow size-dependent electrophoretic mobility.

Bacterial subtyping is the main PFGE application due to the discriminatory power, simplicity and low cost of the technique. Although some laboratories are transitioning towards whole genome sequencing-based typing, it remains important for small hospitals and laboratories with limited resources and it will be the more feasible option for bacterial subtyping by longer (2)(3). This technology plays a key role in modern genomics, as it allows manipulations with DNA of whole chromosomes or their large fragments (4) as is the case, for example, of Cas9-assisted targeting of chromosome segments [CATCH] (5) or DNA fragment size assessment for Next-Gen Sequencing (6).

Biochemistry is an area where highly sophisticated equipment is required. In general, few equipment - i.e. PCR thermocyclers (7) - have an open source version, limiting the development of improved/custom equipment, and the costs of such equipment tended to be very high, which imposes a high barrier for democratizing its usage in many academic labs. PFGE equipment is expensive and exclusively sold by two providers [Bio Rad, Analytik Jena], they cost around USD$30,000 and do not have an open source version. Commercial PFGE equipment control of the electric field reorientation are based on both electronic circuitry [CHEF, Bio Rad] and physical rotating electrodes [Rotaphor, Analytik Jena].

Open hardware and free and open source software [FOSS] provides tools for fabricating devices at lower costs and higher adaptability. Easy to use microcontrollers such as Arduino and manufacturing technologies such as 3D printers has enabled a diverse community of developers to start building new customized, low cost and open source devices.

## 2. Hardware description

### Hardware and circuit design and construction

#### General outline

Here we describe openPFGE, an open source and low cost PFGE equipment. The equipment implements a rotating gel electrophoresis [RGE] over an uniform electric field [Figure 1 & 2], using the basis of the system described by Southern in 1987 (8).The design considers the use of a standard commercially available electrophoresis chamber and the construction of an agarose gel rotation system using 3D printed parts, a servo motor and simple electronics. It incorporates a custom cooling system using a small pump and peltier cooler elements. The control of the motor, the cooling system and the parameters of the run are made via bluetooth communication to a smartphone running an Android app.

**Fig 1.**
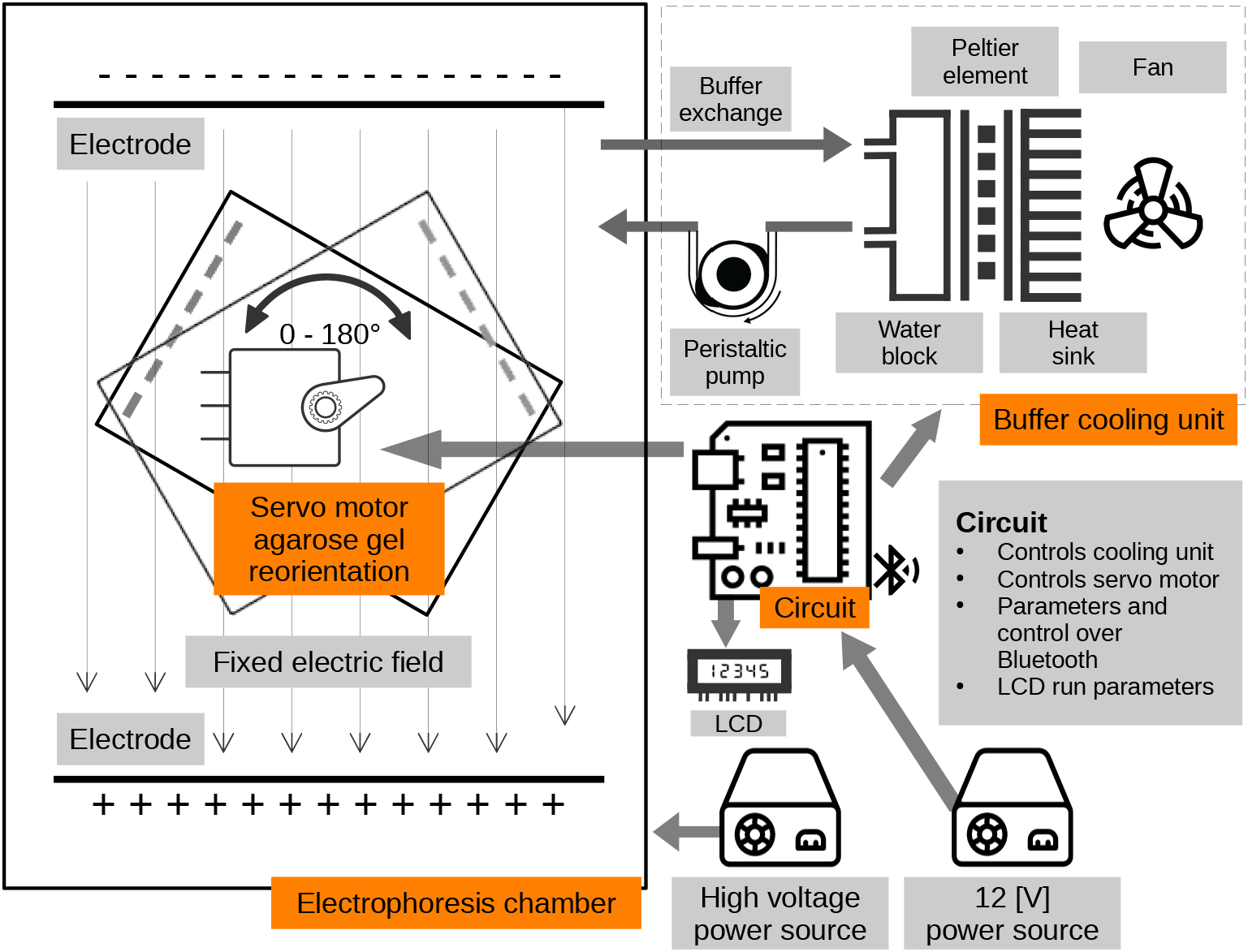
openPFGE equipment diagram. Main components of openPFGE hardware: A servo motor reorient the agarose gel position by rotating a gel tray in respect to a fixed electric field. Movement of the servo motor is controlled by a circuit whose parameters are determined via Bluetooth by an Android app. Circuit also controls a buffer cooling unit.*. *Some icons made by Freepick, Google, Eucalyp and Those Icons from www.flaticon.com.

**Fig 2.**
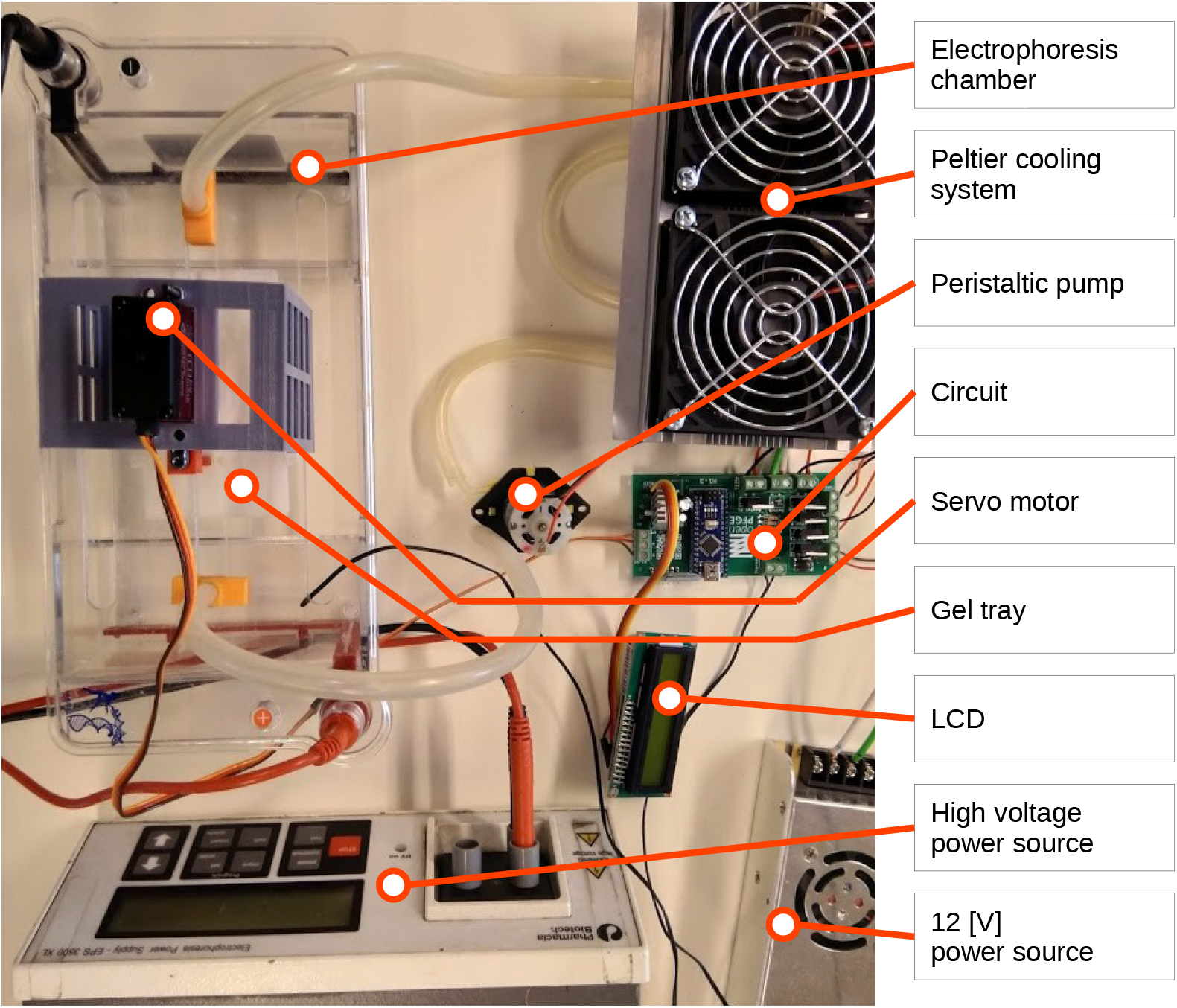
openPFGE equipment picture. General view of the RGE system implemented, highlighting its main components.

#### RGE system

In a RGE, the electrodes of the electrophoresis chamber remains fixed while the agarose gel is rotated to achieve the reorientation of the direction of the electric field across the gel. As the servo motor that performs the movement of the agarose gel in this implementation operates in 180° angle, the position of the gel in respect to the electric field can be specified at any time of the run in a −90° / +90° position. A zig-zag pattern in the direction of the electric field is applied to obtain a net straight migration of DNA molecules [Figure 3].

**Fig 3.**
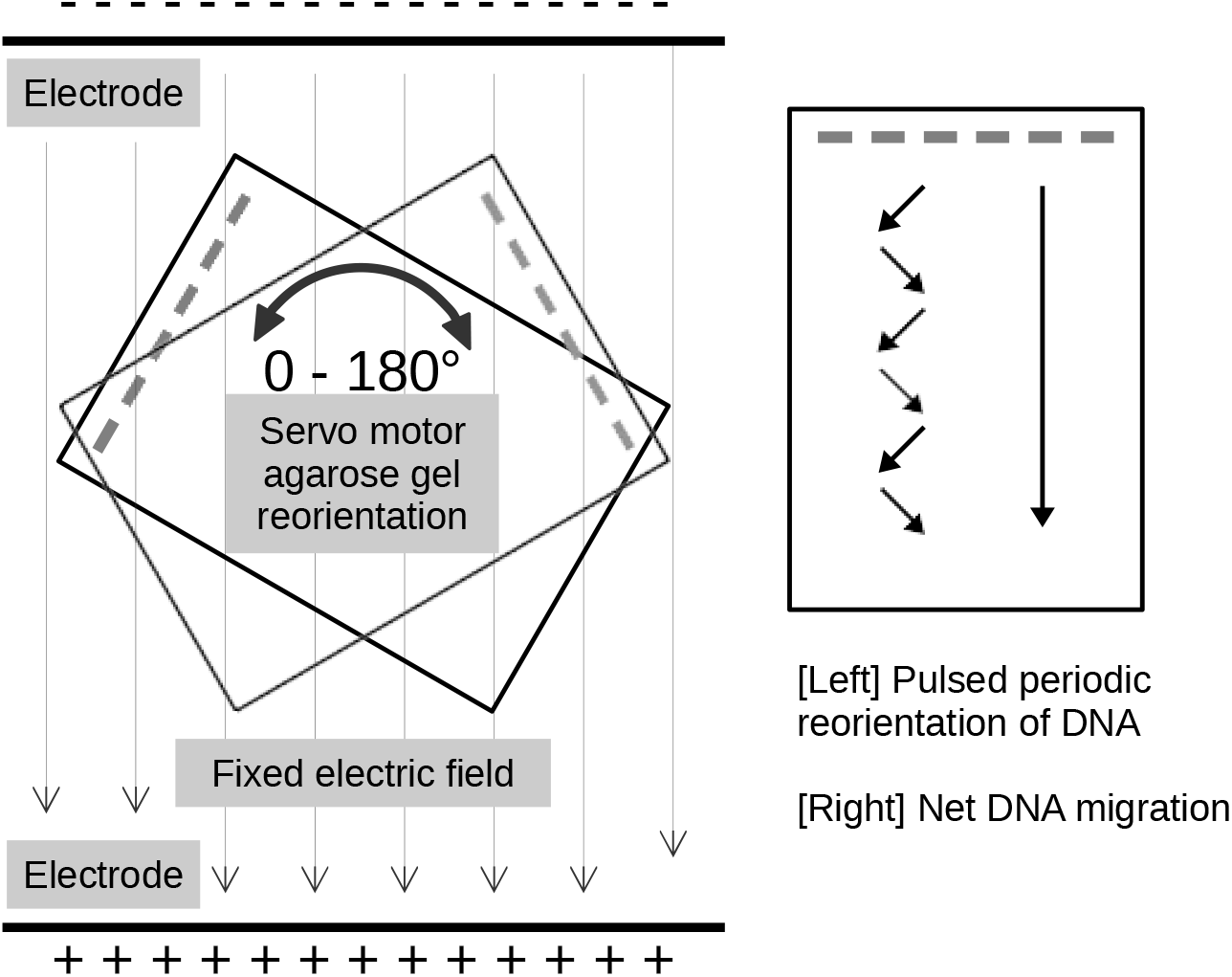
Electric field reorientation in RGE. Diagram shows [Left] the static electric field provided by the electrophoresis chamber electrodes and how the rotation of the gel by the servo motor alters the orientation of the electric field through the agarose gel and [Right] a zig-zag pattern in the direction of the electric field is applied to obtain a net straight migration of DNA molecules

In order to keep the design simple and the cost as low as possible, a gel rotating system inside a standard large electrophoresis chamber was implemented in this design. In this case, a 310 × 150 × 90 [mm] chamber and a 100 × 60 [mm] agarose gel is proposed, but it can be adapted to any chamber/gel size.

This gel rotating system incorporates simple 3D printed parts composed of: 1) a tray base, that supports the gel; 2) a tray cover, that fixes the gel position; 3) a stem, that goes from the tray cover through the electrophoresis chamber cover and joins the servo motor; and, 4) a joint, of the stem to the servo motor that allows easy release of the tray using a strong magnetic couple. It also includes a 180 degrees, 5 [V], high speed and accuracy digital servo motor - like the DS3218-MG - that allows the PFGE to move at any specified angle. All 3D printed parts were designed considering the Fused Deposition Modeling [FDM] process by using mostly horizontal and vertical faces, low inclination angles and no - or really small - ceilings, therefore, no support structures are needed. Figure 4 shows the gel tray and servo joint design.

**Fig 4.**
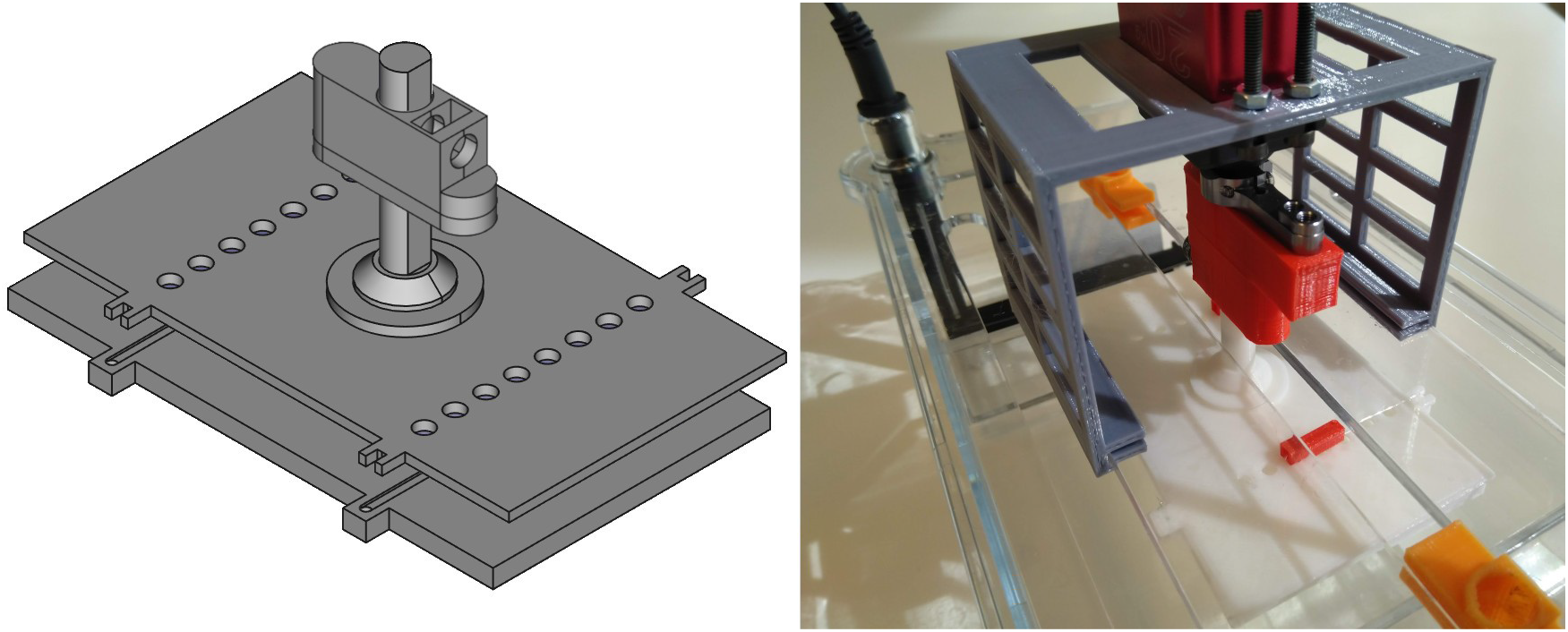
Gel rotating system. [Left] Scheme of the gel tray composed of the tray base, tray cover, stem and tray/servo motor joint and [Right] Picture of the gel tray (white parts) inside the electrophoresis chamber coupled to the servo motor (red parts) standing on the motor support (gray part) that attaches to the chamber cover

#### Cooling system

The cooling system is composed of a small 12 [V] peristaltic pump that circulates the buffer through a peltier cooler element based refrigeration system consisting of two peltier 12 [V] / 5 [A] elements, an aluminum heat dispenser and two 12 [V] fans [Figure 5]. An epoxy covered NTC thermistor allows to sense the current buffer temperature in order to define the start/stop of the cooling system.

**Fig 5.**
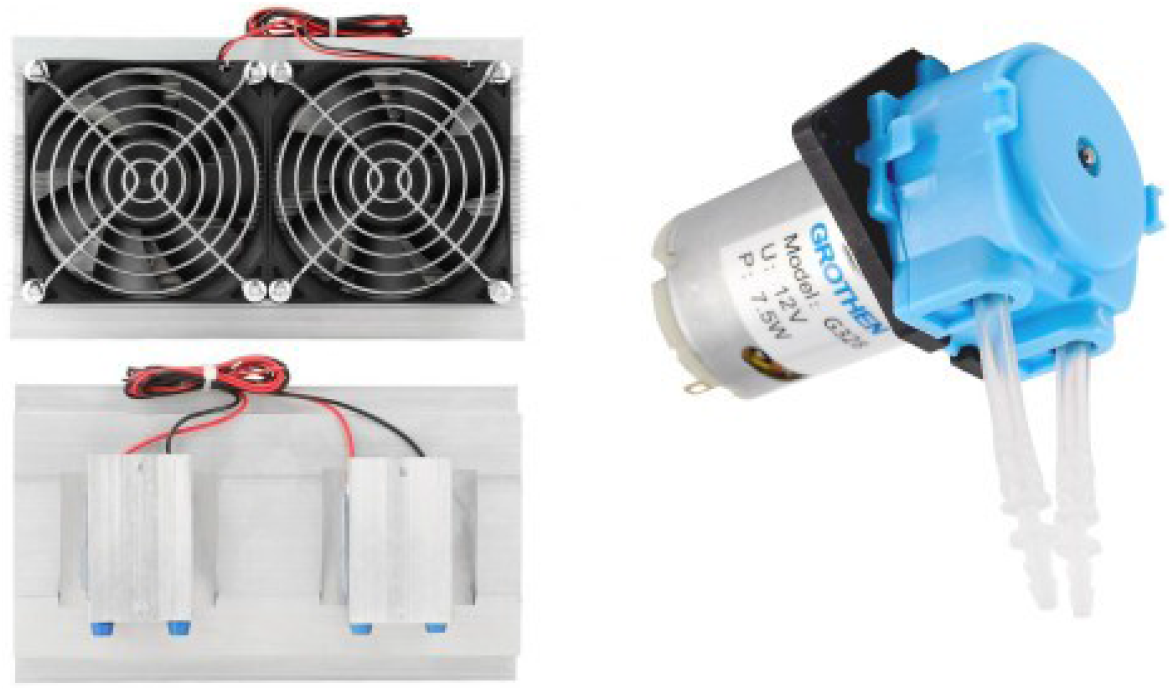
Cooling system components. [Left] Peltier cooler system composed of peltier elements, fans and water aluminum block, top and bottom views and [Right] Peristaltic water pump

#### Electronic components

The circuit is driven by an Arduino Nano microprocessor. It is supplied by a 12 [V] / 30 [A] power source and has a 5 [V] regulator (LM7805) to provide power to the microprocessor, components and servo motor. It uses a HC-05 Bluetooth module to provide communication to a smartphone for system control. It holds a serial LCD/I2C module to display the parameters of the electrophoresis run independent from the smartphone. The NTC thermistor signal is read by an analog input and peltier elements, fans and the pump are controlled by digital outputs through high current MOSFETs. The PCB design can be found at the Gitlab repository (9) or Zenodo repository (13).

The electrodes of the electrophoresis chamber are connected to a standard commercial electrophoresis power source. It has to be manually operated as the circuit of the openPFGE does not have, yet, control control over its function.

### Software

Agarose gel positioning and buffer temperature control are the main variables controlled by the software. The servo motor moves the agarose gel in a maximum range of 180°, −90° / +90° in respect to the direction of the fixed electrophoresis electric field, in a zig-zag pattern [Figure 3]. Some PFGE protocols require constant time on each position of the electric field at each step while other requires time ramps. In time ramps, the protocol defines a initial time of the electric field at each step at the beginning of the run (ramp start time), a final time (ramp end time) and a total run time, so incremental step times from ramp start time to ramp end time trough total run time is executed. The software performs the calculation of the ramp time at each step based on these three parameters.

Angle is an important factor in achieving optimal resolution. Separations of larger sizes of DNA are greatly improved by the use of a smaller angles. Using a smaller results in savings in run time (up to 50%) (14). openPFGE receives fixed angle as a parameter of the run and can take values from 0° to 180°.

The cooling system is activated upon demand to achieve the buffer temperature setpoint provided by the user. It turns fans/peltier elements pairs on/off according to the offset from setpoint value.

#### Firmware

The Arduino Nano microprocessor runs a firmware that allows the control of the motor rotation and refrigeration system, the read of analog signal from the thermistor and the communication of the current parameters to the LCD/I2C module display and by bluetooth with the Android app that gives the parameters of the run. It allows the run to have a fixed rotation time or a growing rotation time ramp. Time ramp consists of three parameters: start run time, end run time and run duration. The firmware allows the equipment to run normal PFGE. Other run modes (as FIGE or CHEF) are expected to be available in the next firmware/app update.

#### Android app

A dedicated android app was developed in order to control the equipment. It establishes communication with the PFGE via Bluetooth and capture user preferences as: on/off, pause, ramp on/off, ramp start time, ramp end time, ramp duration time, wait on position time (fixed step time), rotation angle, total run time, buffer temperature (setpoint), temperature automatic control on/off, among other options. Is has the option of load predefined run protocols for commercial markers as well to save user defined programs. Figure 6 shows the main views of the app. App download is available at Google play store (10).

**Fig 6.**
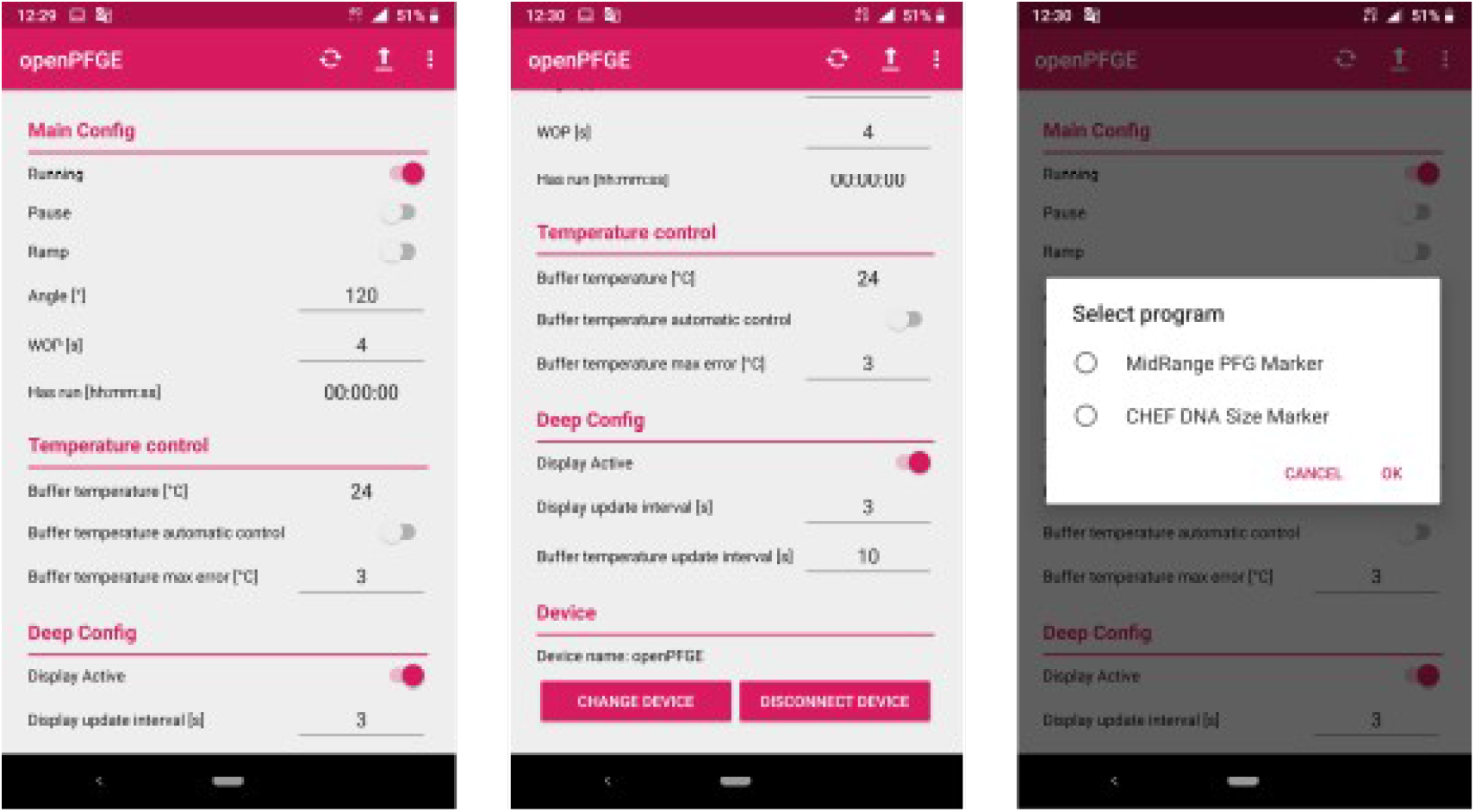
openPFGE app main views. *Left*: Main view of the app showing the control and monitor options. *Center*: Continuation of the main view. *Right*: View that shows how to load predefined and custom run protocols

### Repository and open source hardware certification

All 3D parts design files, code and electric diagrams can be found at the Gitlab repository (9) or Zenodo repository (13).

The equipment has been certified under OSHWA certification: [OSHW] CL000001 | Certified open source hardware | oshwa.org/cert.

### Costs

Table in *Bill of materials* section summarizes the costs of each component of the equipment, including electrophoresis chamber and power source. Commercial PFGE equipment is around USD$30,000. The total cost of openPFGE equipment is about USD$850, therefore it is about 3% of the price of a typical commercial equipment, considering electrophoresis power source and chamber.

## Discussion

In order to contribute to the democratization and acceleration of hardware development in biochemistry, we present in this work a certified open source hardware that implements rotating gel electrophoresis to perform PFGE. All 3D parts design files, code and electric diagrams are open source and publicly available at the Gitlab repository (9) or Zenodo repository (13). The equipment is capable of the separation of DNA molecules up to, at least, ~2 Mbp, costs about USD$850, about 3% of the price of typical commercial equipment considering the chamber and the power source. OpenPFGE has similar requirements, capabilities and results than commercial equipment in terms of applications, maximum fragment size resolution and experiment run time/setup. As it is the first version of the equipment, further improves in design of components, electronics and software can lead to a more robust implementation. This, together with validation of building and use by other research groups could end in a equipment with similar application confidence than commercial equipment. Although open blueprints for PFGE instrument has been described earlier (11), to our knowledge, this is the first PFGE fully documented, ready to build, open source, flexible and modern equipment available.

Other modes of use such as field inversion gel electrophoresis [FIGE] are intended to be added in the future among other applications like, for example, DNA size selection for Next-Gen Sequencing. Open source versions of a gel electrophoresis chamber (16) and electrophoresis power supply (17) are publicly available. Is expected to develop versions of these components to fit openPFGE in order to bring a whole open source PFGE equipment, with both lower costs and flexible design.

The openPFGE

- Allows for run standard PFGE at low equipment cost
- Allows for easy and inexpensive maintenance and repair of the PFGE equipment
- Establishes the first open source PFGE development that enables future equipment improvement by the community

## Materials and methods

### 3D printed parts

All 3D parts were designed using Freecad software and printed using PLA filament and Ender 3 or 5 printing machines at Fablab U. de Chile (12). Parts designs and detailed printing conditions can be found at the Gitlab repository (9) or Zenodo repository (13).

### Electronic components and PCB design

PCB design was made using Fritzing software. All electronic components and PCB design can be found at the Gitlab repository (9) or Zenodo repository (13). PCB (H1.3) is ready to fabricate on Aisler company (15).

### Software

Arduino IDE software was utilized for the microcontroller programming. Android studio was utilized for the app development, using API 29 (Android 10) and min API 26 (Android 8).

## 3. Design files

### Design Files Summary

| Design file name | File type | Open source license | Location of the file |
| --- | --- | --- | --- |
| *motor joint.FCStd* | CAD | CERN | https://zenodo.org/record/3727331 |
| *motor support.FCStd* | CAD | CERN | https://zenodo.org/record/3727331 |
| *pin.FCStd* | CAD | CERN | https://zenodo.org/record/3727331 |
| *tray.FCStd* | CAD | CERN | https://zenodo.org/record/3727331 |
| *tray cover.FCStd* | CAD | CERN | https://zenodo.org/record/3727331 |
| *tray cover joint.FCStd* | CAD | CERN | https://zenodo.org/record/3727331 |

The use of each of the design files is best determined by the descriptions and rendering shown in *Build instructions* section.

## 4. Bill of Materials

Costs to build 1 unit of an openPFGE equipment, including electrophoresis power source and chamber

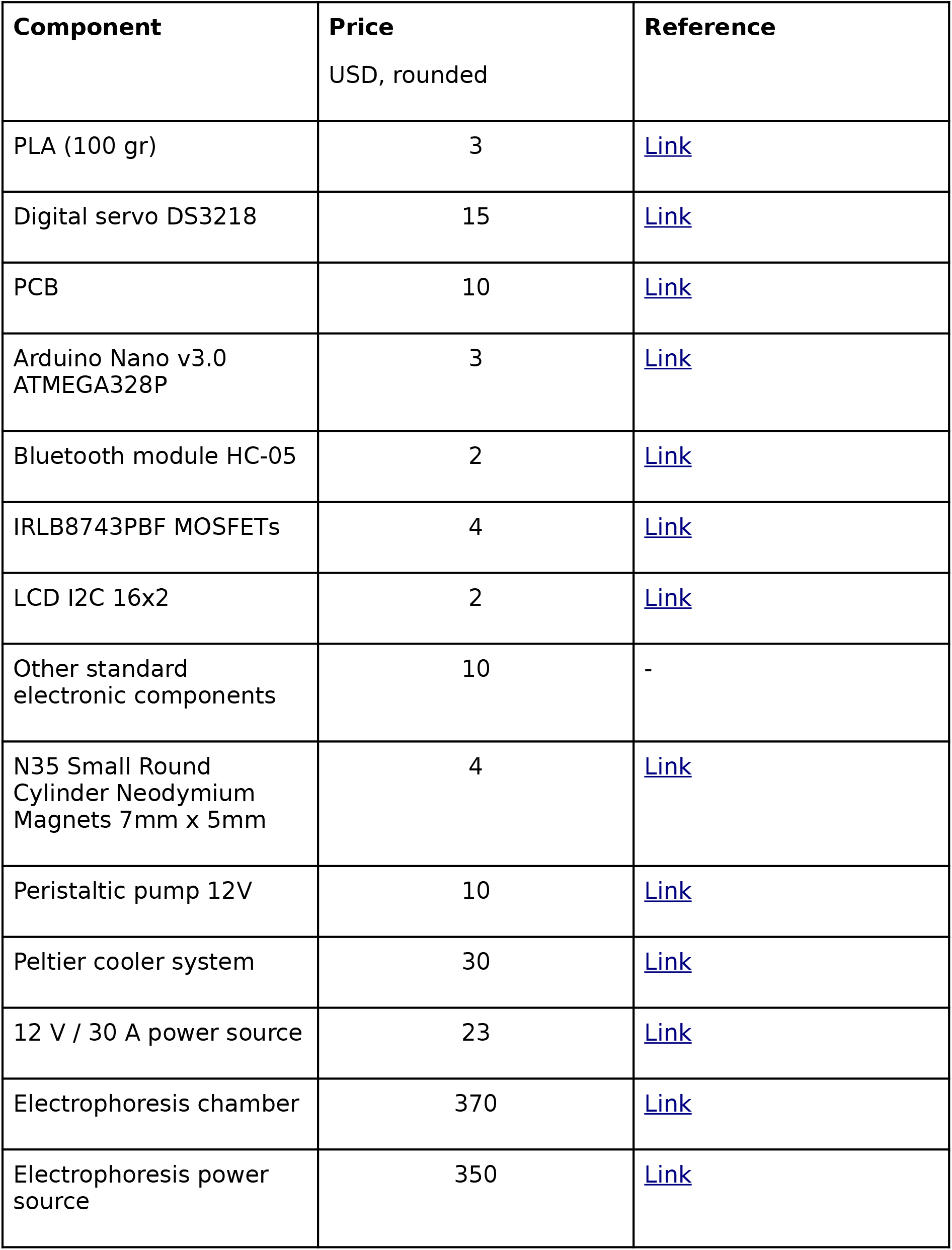

| **Component** | **Price**<br>USD, rounded | **Reference** |
| --- | --- | --- |
| PLA (100 gr) | 3 | Link |
| Digital servo DS3218 | 15 | Link |
| PCB | 10 | Link |
| Arduino Nano v3.0<br>ATMEGA328P | 3 | Link |
| Bluetooth module HC-05 | 2 | Link |
| IRLB8743PBF MOSFETs | 4 | Link |
| LCD I2C 16x2 | 2 | Link |
| Other standard<br>electronic components | 10 | - |
| N35 Small Round<br>Cylinder Neodymium<br>Magnets 7mm x 5mm | 4 | Link |
| Peristaltic pump 12V | 10 | Link |
| Peltier cooler system | 30 | Link |
| 12 V / 30 A power source | 23 | Link |
| Electrophoresis chamber | 370 | Link |
| Electrophoresis power<br>source | 350 | Link |

## 5. Build Instructions

A. **PCB and electronics**. All electronic parts are listed in the *readme* file on the Gitlab repository (9) or the Zenodo repository (13)

1. **Manufacturate your PCB**: using the *Circuit/openPFGE H*.*.fzz* file. Version H1.3 is ready to be ordered at Aisler company (15).
2. **Solder the components**: The PCB skilscreen shows exactly the position of each component. The components that should be soldered are:

✓ Arduino Nano v3.0 ATMEGA328P
✓ Bluetooth module HC-05
✓ L7805CV Transistor voltage regulator (1.5 A) with a heat dissipator (not shown in the picture below)
✓ 100 uF capacitor
✓ 10 uF capacitor
✓ 4 × 1N4007 Diode
✓ 6 × 10K resistor
✓ Power switch (not shown in the picture below)
✓ LCD I2C 16×2 (just cable connections shown in the picture below)
✓ 8 × 2 Pin Plug-in Screw Terminal Block Connector
✓ 1 × 3 Pin Plug-in Screw Terminal Block Connector

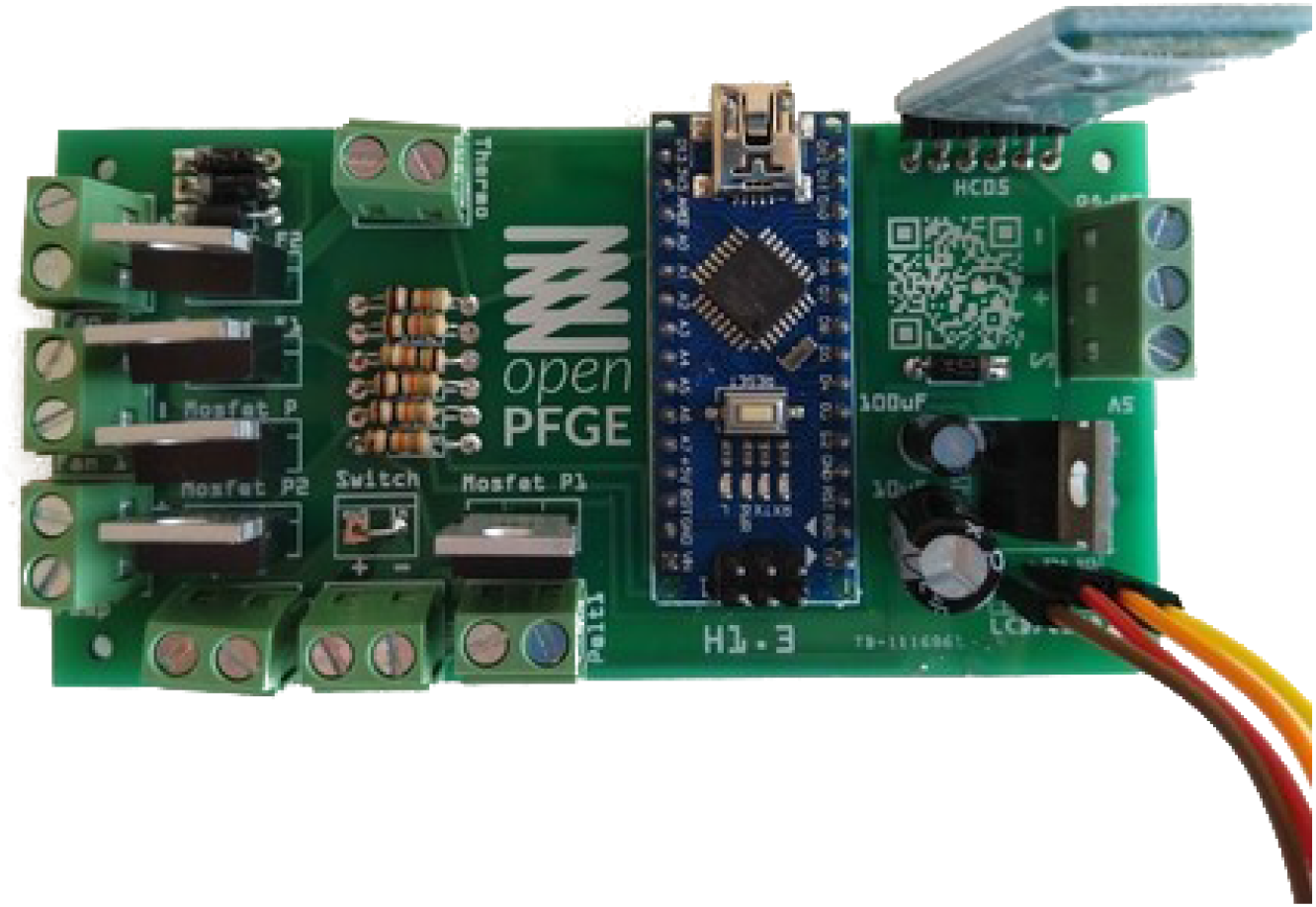
*Final PCB with all its components soldered*
3. **Connect the peripherals**: The components connected using the terminal blocks are:

✓ Epoxy NTC thermistor temperature sensor 10K 1%
✓ 12V / 30 A power source
✓ Digital servo DS3218
✓ Peristaltic Pump 12V
✓ Peltier cooler 12V

♦ 2 fans
♦ 2 peltier elements
4. **Upload the firmware**: Using Arduino software:

✓ On *Tools* > *Board* select *Arduino Nano*
✓ On *Tools* > *Processor* select *ATMEGA328P (Old bootloader)*
✓ [optional] Set the name of the Bluetooth module by uploading the code on *set_bt_name_password.ino* file and by sending AT commands [follow this instructions]
✓ Confirm you have all the required libraries loaded on your Arduino according to the includes in the firsts lines of code in the *openPFGE_F_X_X.ino* file
✓ Compile and upload the code in the *openPFGE_F_X_X.ino* file to the Arduino Nano module using an USB cable
B. **3D parts**. This parts allows to mount the motor in the electrophoresis chamber cover and to build the gel tray

1. **Motor support**

✓ Print the *motor support.FCStd* and the *motor joint.FCStd* files using PLA or ABS materials
✓ Mount the servo motor to the support using 4 screws
✓ Couple the joint to the servo motor arm using a screw in the axis and other in the extreme of the arm
✓ Paste, using glue, two round 7mm × 5mm magnets to the motor joint in the remaining holes
✓ Mount the motor support to the cover of the electrophoresis chamber

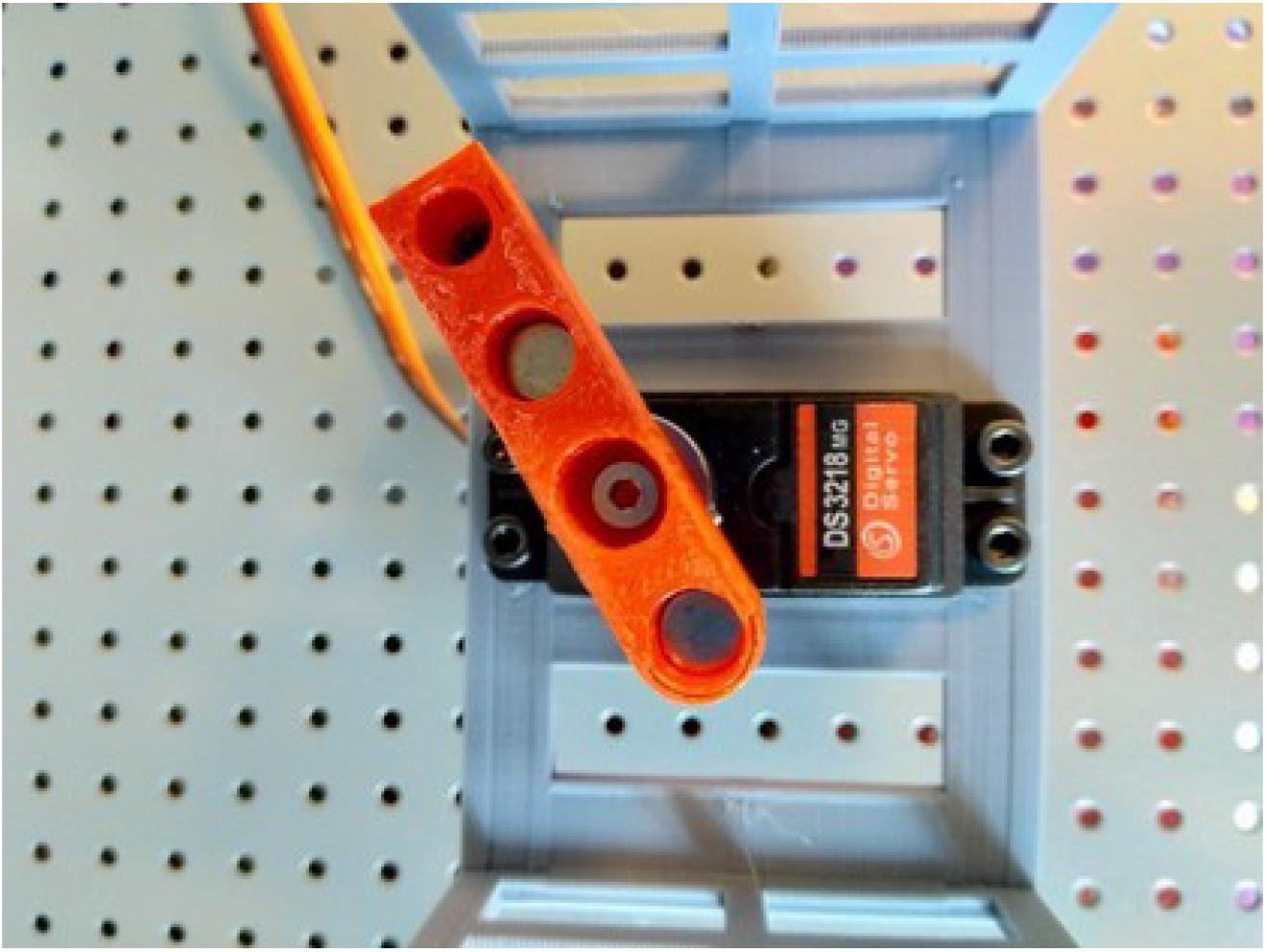
*Motor mounting*
2. **Gel tray**

✓ Print the *tray.FCStd*, *tray cover.FCStd*, *tray cover joint.FCStd* and 2 × *pin.FCStd* files using PLA or ABS materials
✓ Paste using glue two round 7mm × 5mm magnets to the tray cover joint and add a screw-nut pair that fixes the piece to the stem of the tray cover
✓ Use some fishing thread to join from the tray cover, through the tray (base), up to the pins in order to tie up the tray cover and the tray (base)
3. **Hose adapters**

✓ Print two *hose_adapter.SLDPRT* files using PLA or ABS material

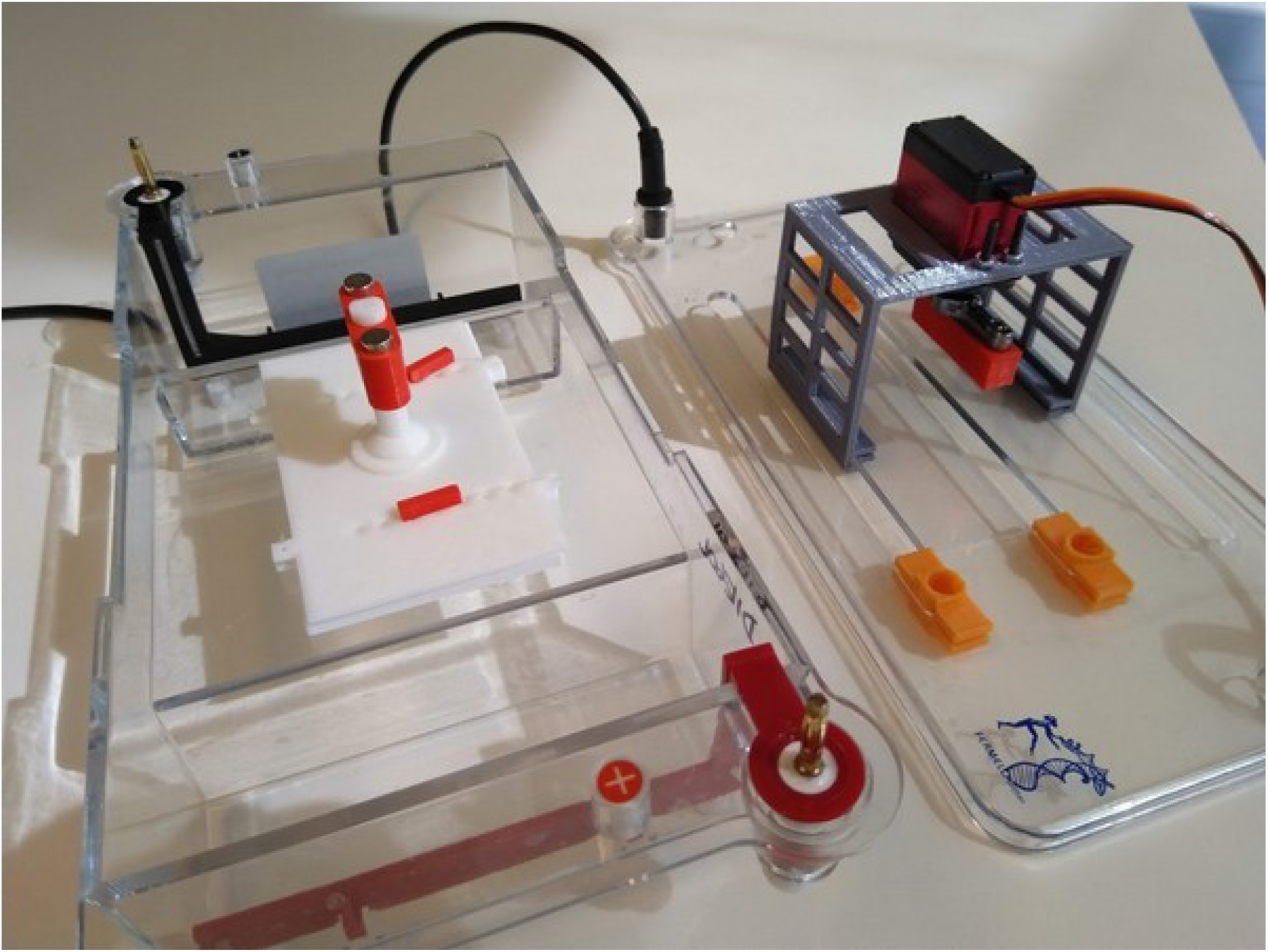
*Final assembly of the printed parts in the electrophoresis chamber*
C. **Assembly**. Connect the parts

1. **Cooling unit**

✓ Circuit from the first hose adapter in the electrophoresis chamber cover through the peristaltic pump, the water blocks of the peltier cooling system and back to the second hose adapter using an appropriated hosepipe
✓ Make sure the temperature sensor, pump, peltier elements and fans are connected to the circuit as indicated in part A
2. **Gel tray, chamber and servo motor**

✓ Put the gel tray (base and cover) inside the electrophoresis chamber
✓ Adjust the vertical position of the tray joint by using the screw that fixes the piece to the stem of the tray cover to fit the counter-part in the servo motor
✓ Place the cover of the electrophoresis chamber and join the gel tray and the servo motor using the magnetic joint
✓ Make sure the temperature sensor and the servo motor are properly connected to the circuit as indicated in part A
3. **Ready to run**

✓ Connect the electrodes to the high voltage power source
✓ Power on the high voltage power source and the 12[V] power source that provides energy to the circuitry
✓ Use the android app to set the run parameters and control the start of the run
✓ See “operation instructions”

## 6. Operation Instructions

### Safety concerns

1. **Power source:** The 12 [V] power source input is connected to high voltage. Make sure this connection is properly isolated and without risks of water/buffer dropping. Be sure the power source has enough fresh circulating air in order to eliminate the heat produced by the circuit.
2. **Circuit**: The circuit is powered by 12 [V]. This may not be considered dangerous for the operator. Make sure this circuit is properly isolated and without risks of water/buffer dropping in order to prevent circuit damage. Be sure the circuit has enough fresh circulating air in order to eliminate the heat produced by the circuit.

### To make a PFGE run

1. Download and install the *openPFGE* Android app from Google Play Store (10)
2. Load your agarose gel with the samples on the gel tray. Close the gel tray using the *pins*
3. Introduce the tray to the electrophoresis chamber and connect the tray to the motor join and close the electrophoresis chamber. Make sure to level the gel tray about 1[cm] above the bottom of the chamber using the screw provided for this.
4. Add the appropriated electrophoresis buffer to exceed the top of the gel tray in about 1[cm]
5. Power on the 12V transformer of the circuit
6. Use the smartphone menu to pair Blue tooth to the openPFGE-*** module
7. Open the openPFGE app and configure the run parameters
8. Tap the send button on the menu bar and wait for the confirmation of *successful received from device* message
9. Power on and start the electrophoresis power source. That’s all

## 7. Validation and Characterization

### PFGE marker validation

Two commercial available markers were tested using openPFGE equipment under similar run conditions as the provider recommendations in terms of buffer, agarose concentration, ramp times and rotation angle. Figure 7 and Figure 8 shows the comparison between a commercial marker reference image and the result obtained with this equipment for the MidRange PFG Marker (Max size band ~250kbp) and CHEF DNA Size Marker #1703605 (Max size band ~2.2Mbp).

**Fig 7.**
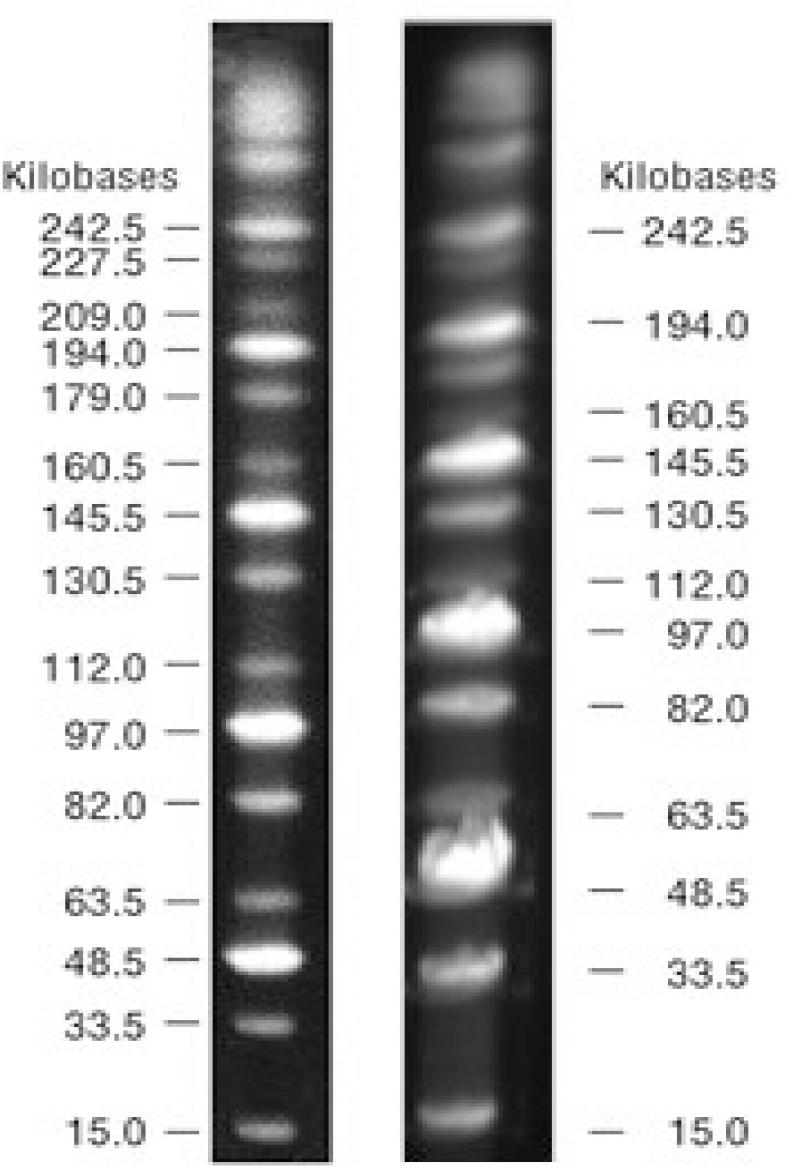
250 kbp marker comparison. MidRange PFG Marker (*left*) provider reference image and (*right*) run in openPFGE equipment

**Fig 8.**
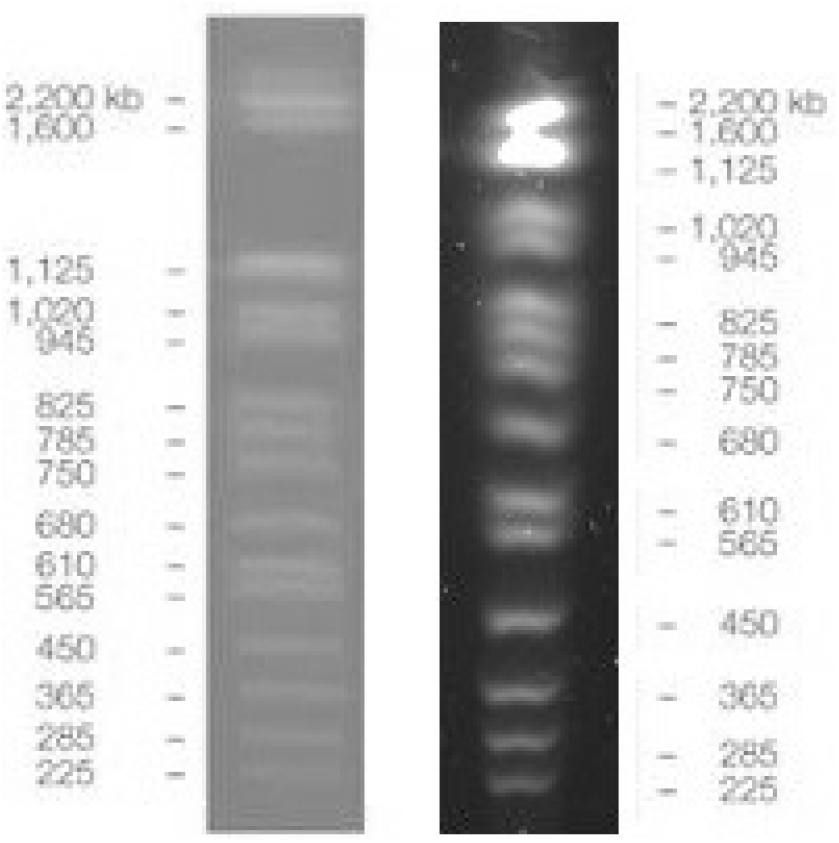
~2.2Mbp marker comparison. CHEF DNA Size Marker #1703605 (*left*) provider reference image and (*right*) run in openPFGE equipment

### Materials and methods

#### PFGE markers

MidRange PFG Marker (N0342S) was purchased from New England Biolabs. CHEF DNA Size Marker, 0.2–2.2 Mb, S. cerevisiae Ladder (1703605) was purchased from Bio-Rad Laboratories, Inc.

#### PFGE marker validation

In order to validate the operation of the equipment, two commercial markers were tested under the following conditions, using a PowerPac 200, Bio Rad, electrophoresis power supply: 1) MidRange PFG Marker: 0.5X TBE, 6 volts/cm, angle 120 [°], ramp start time 1 [s], ramp end time 25 [s], run time 12 [h], buffer temperature 12 [°C]; and 2) CHEF DNA Size Marker #1703605: 0.5X TBE, 6 volts/cm, angle 120 [°], ramp start time 60 [s], ramp end time 120 [s], run time 16 [h], buffer temperature 12 [°C].

## 9. Credit Author Statement

**Diego Lagos-Susaeta**: Conceptualization, Methodology, Software, Validation, Investigation, Resources, Writing - Original Draft, Visualization. **Oriana Salazar**: Supervision. **Juan A. Asenjo**: Supervision.

## 10. Acknowledgements

We would like to thank the equipment and advisement provided by staff and developers at Fablab U. de Chile (12).

